# DiaReport: Reproducible Workflow for Differential Expression Analysis and Interactive Reporting in DIA-based Proteomics

**DOI:** 10.64898/2026.03.10.707945

**Authors:** Andrea Argentini, Esperanza Fernández, Jarne Pauwels, Kris Gevaert

## Abstract

**Summary:** Data-independent acquisition (DIA) has become the preferred data acquisition method for mass spectrometry-based proteomics, yet, reproducible workflows for differential expression (DE) analysis and reporting results remain limited. We present DiaReport, an R package that performs precursor- and protein-level DE analysis from DIA-NN output using MSqRob and QFeatures, while generating high-quality, interactive HTML reports through Quarto. DiaReport integrates precursor data, filtering of missing values, normalization, protein summarization and statistical modeling within a single function, supporting both simple pairwise as well as complex experimental designs. The package provides structured outputs and configuration files to ensure computational reproducibility across different studies. To accommodate diverse research needs, DiaReport includes multiple reporting templates tailored to different proteomic applications. Applying DiaReport to an extracellular vesicle (EV) proteomics dataset demonstrates its ability to efficiently analyze DIA data and provide rapid insights into sample quality and protein level differences.

**Availability and Implementation:** DiaReport is an open-source R package available at https://github.com/Gevaert-Lab/diareport. The package is platform-independent and distributed under the MIT license. Reports are generated using Quarto and require only standard R dependencies. Detailed documentation, installation guides and usage vignettes are provided within the repository. The interactive HTML reports discussed in this study, including the UPS2 benchmark and EV case study, are archived on Zenodo (https://doi.org/10.5281/zenodo.18632744 and https://doi.org/10.5281/zenodo.18632731).

**Contact:** Corresponding author: Kris Gevaert.

## Introduction

In mass spectrometry-based proteomics, data-independent acquisition (DIA) is rapidly gaining prominence over data-dependent acquisition (DDA)^1,2^. Unlike DDA, DIA systematically fragments all peptide ions within a defined mass-over-charge range, increasing the number of identified peptides and enabling more robust quantification. Among the available open-source tools for DIA data processing, DIA-NN^3^ has emerged as a standard tool, supported by frequent updates and an active user community^4^.

Differential expression (DE) analysis is a routine step in quantitative proteomics. Several R packages support this task, including MSstats^5^, limma^6^ and MSqRob^7,8^. The MSstats ecosystem provides modality-specific workflows for Tandem Mass Tag (TMT) labeling, Label-Free Quantification (LFQ), post-translational modifications (PTMs)^9^ and Limited Proteolysis coupled to Mass Spectrometry (LiP-MS)^10^, and supports multiple data sources, including DIA-NN output. MSqRob, particularly when paired with QFeatures^11^, provides a flexible statistical framework capable of handling complex experimental designs, as demonstrated in PTM^12^ and TMT workflows^13^. However, despite the availability of strong individual components (e.g., DIA-NN for quantification and MSqRob for modelling), few tools integrate these steps into a reproducible and automated end-to-end workflow with high-quality reporting. One example addressing parts of this problem is prolfquapp^14^, a command-line tool that generates differential expression analysis reports based on the prolfqua package^15^ and that provides an R Shiny application for interactive visualization of differential expression results and gene set enrichment analysis (GSEA). In contrast, DiaReport integrates data processing, statistical modeling and high-level reporting within a single, streamlined R package. This eliminates the need for switching between command-line interfaces and does not rely on external Shiny applications for visualization as DiaReport generates self-contained, portable HTML reports. This ensures that the visualizations remain linked to the specific analysis version and can be easily shared or archived without requiring an active server environment.

Reporting is a critical component of data analysis, enabling users to explore results interactively and contextualize biological findings. Frameworks such as Quarto^16^ greatly simplify the creation of interactive HTML reports with Plotly-based visualizations^17^ and dynamic tables^18^. While interactive reporting tools exist for quality control in MS-based proteomics^19–21^, equivalent frameworks for DE analysis remain limited. Computational reproducibility is another essential aspect of modern proteomic workflows. Achieving reproducible analysis requires transparent documentation of processing steps, consistent software environments and standardized output formats^22^. Existing R-based proteomic workflows often rely on ad hoc scripts or loosely structured objects, making analyses difficult to reproduce while limiting interoperability with downstream tools. As DIA datasets grow in scale and complexity, such limitations increasingly hinder routine quantitative proteomics.

Here, we present DiaReport, an R package that performs precursor-and protein-level DE analysis of DIA-NN output using MSqRob and generates fully interactive HTML reports for DE analysis powered by Quarto. DiaReport provides a single entry-point function to execute the entire workflow, automatically applies the selected analysis template and stores results and configuration files in a structured directory format. This design enhances reproducibility and was created to simplify collaboration and facilitate deployment within research groups with limited programming or IT expertise.

### Implementation

The DiaReport R package consists of two main components: (i) a data-processing module that prepares DIA-NN quantitative features for differential expression analysis and (ii) a reporting module that generates an interactive HTML report using Quarto templates (Figure 1A). Both precursor-and protein-level workflows are supported within a single unified function interface.

**Figure 1.**
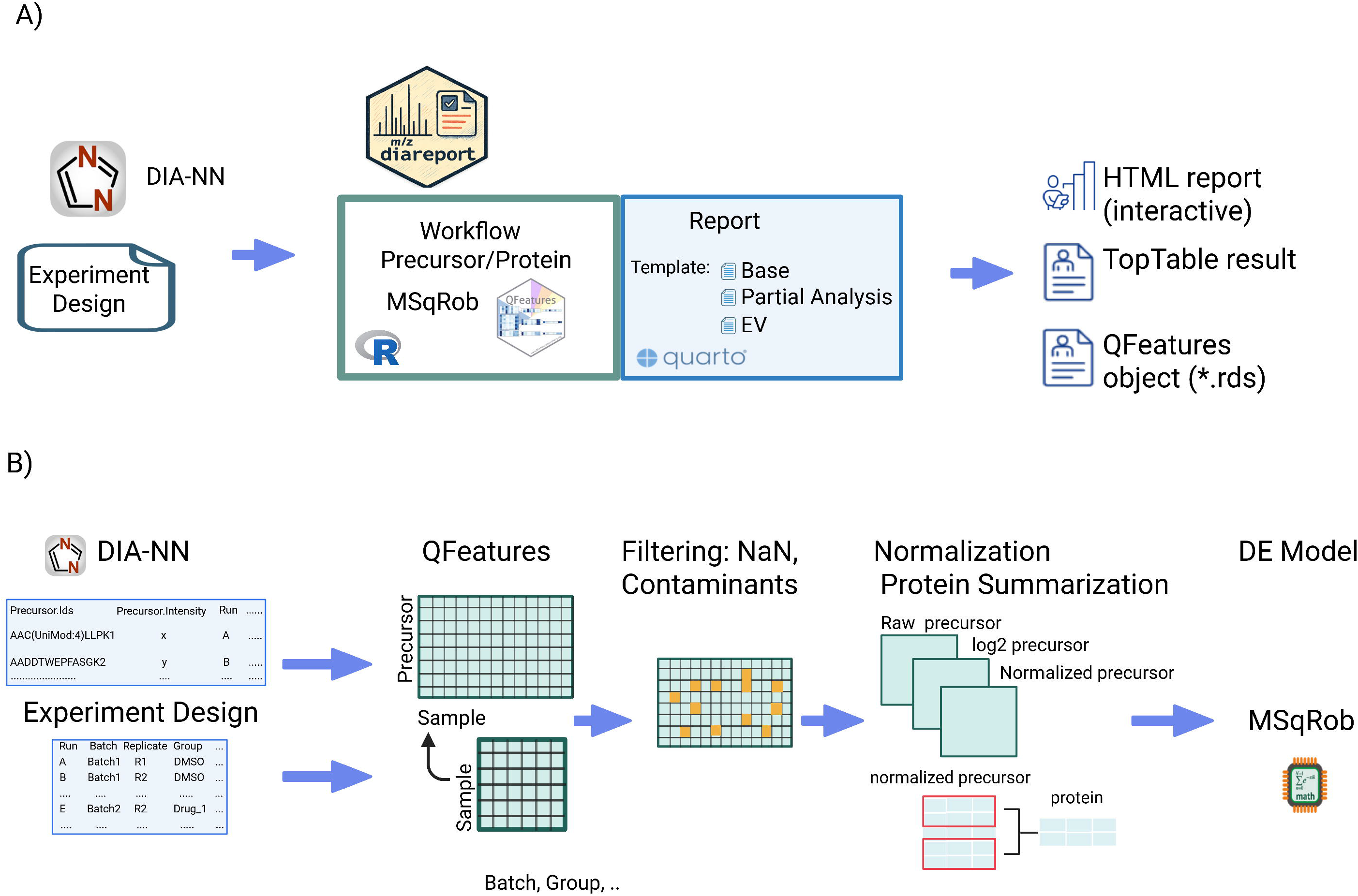
Overview of the DiaReport package and workflow. (A) Package architecture: DiaReport streamlines DIA-NN data processing using MSqRob and QFeatures, generating structured outputs including interactive HTML reports, TopTable results and a serialized QFeatures object for downstream analysis. (B) Precursor-level data and experimental metadata are integrated into a QFeatures object. The pipeline automates filtering (missing values, contaminants), log_2_-transformation and normalization before precursor-to-protein summarization and statistical modeling with MSqRob.

### Experiment design file (EDF)

DiaReport requires two input files: DIA-NN report output (as parquet file or tsv for DIA-NN version < 1.8) and an experiment design file (EDF). The latter is a comma-separated table that specifies sample metadata (sample name, raw file, replicate and experimental group). Optional variables, including covariates and confounders, can be incorporated into the statistical model.

### Data processing and differential analysis

The workflow follows a sequential pipeline: (i) precursor-level q-value and contaminant filtering, (ii) completeness-based filtering using the user-defined k threshold, (iii) log_2_-transformation and normalization and (iv) robust protein-level summarization (Figure 1B). Precursors that pass the fixed DIA-NN precursor-and protein-level q-value thresholds (q ≤ 0.01) are used to populate a QFeatures object using Precursor.Quantity as the quantitative feature. As MSqRob explicitly models missing values, imputation strategies are not currently applied. To address the inherent prevalence of missing values in proteomics, DiaReport implements three completeness-based filtering strategies, defined by a user-specified completeness threshold k (expressed as a percentage): per-group completeness requires at least k percent valid values within *at least one* experimental group; across-groups completeness requires at least k percent valid values across *all* experimental groups simultaneously; and global completeness requires at least k percent valid values across the entire dataset, regardless of group assignment. These strategies allow users to balance the inclusion of condition-specific proteins (per-group) against the need for high-confidence features with minimal missingness (across-groups/global). Additional filters remove contaminants, restrict analysis to proteotypic precursors and ensure a user-defined minimum number of (different) precursors per protein. Intensities are log_2_-transformed and normalized using user-selected methods. In the protein-level workflow, precursor intensities are summarized using several supported approaches, including median polish and robust summarization (Figure 1B).

Differential expression analysis is performed using MSqRob at either the precursor or protein level. The user specifies the experimental design via an R formula argument (e.g. ∼ condition + batch), allowing for the flexible incorporation of multiple covariates or blocking factors.

### Reporting layer and template

The reporting layer renders interactive HTML reports using Quarto templates. The architecture maintains a strict separation between data analysis logic and visualization, allowing for modularity and future expansion through custom templates. Currently, DiaReport includes three templates designed to meet increasing levels of analytical depth: the base template essentially provides quality control and differential expression analysis; the partial template adds ‘absent-from-DE’ analysis to identify features exclusively detected in specific condition groups; and the EV template offers domain-specific workflows for extracellular vesicle research, including marker panel summaries and contaminant interference analysis, which can be adapted to other biological contexts by supplying custom marker panels. While all templates support pairwise comparisons, the base workflow provides the greatest flexibility, enabling more complex experimental designs such as factorial models (Figure 1B).

To ensure a clean and reproducible environment, processed results are saved as an RDS file and rendered from a temporary directory. All templates share a core set of features, including missing-value diagnostics, completeness summaries and PCA plots. Within the report, each comparison is organized into dedicated tabs containing interactive volcano plots, heatmaps and searchable result tables. The partial and EV templates further extend these capabilities by highlighting condition-specific features, with the EV template providing specialized visualizations for EV-specific quality metrics.

### Structured output

In addition to differential expression analysis and interactive reporting, DiaReport enhances computational reproducibility by storing all analysis parameters in a YAML configuration file and improving result interoperability by saving all analytical outputs within a QFeatures object. Moreover, for each comparison, all plots generated in the report are exported in PDF format and the corresponding top-table results are saved as CSV files within a standardized, structured directory layout. These structured outputs, together with the QFeatures data objects, enable seamless integration with downstream tools, including R Shiny applications for interactive visualization of gene enrichment analyses.

## Results

### Base template analysis

DiaReport was validated using a UPS2/yeast benchmark where the Universal Protein Standard 2 (UPS2) was spiked into a constant yeast background at varying concentrations^4^ (DIA-NN v1.8.1, *in silico* library). The base template accurately recovered expected protein fold-changes, with UPS2 proteins consistently upregulated. Interactive PCA plots showed distinct clustering by spike-in levels, while UpSet plots confirmed high identification overlap across comparisons (see <u>report</u>).

### EV template analysis

To demonstrate DiaReport’s utility on a dataset extending beyond conventional quantitative benchmark studies, the tool was further evaluated on an in-house generated proteomics dataset comparing two different EV enrichment protocols: ultracentrifugation (UC) and an ultrafiltration strategy using 96-well plates (UF96)^23^. Mass spectrometry data were processed using DIA-NN v2.2.0. The <u>EV report</u> generated by DiaReport was specifically tailored to gain rapid and comprehensive insights in the EV proteomics data by plotting several quality metrics, such as the abundance of several EV-specific protein markers (e.g., CD63 and CD81) and interference of contaminants at the precursor intensity level. In addition to EV-specific sections, the report summarizes key analytical outcomes, including the number of features retained after filtering, the effects of normalization and data completeness at the precursor-and protein-level across samples and conditions (Figure 2B) DiaReport analyses revealed a markedly lower abundance of bovine contaminant precursors in the UF96 samples compared with UC samples (Figure 2A), in line with previously reported findings and key for in-depth EV proteome analysis^23^. This was accompanied by a lower variability in EV protein marker abundance across UF96 replicates (Figure 31). Interactive PCA plots revealed a clear separation between the two isolation methods: UF96 triplicates formed a tight, well-defined cluster, whereas UC triplicates showed increased variability, with one replicate deviating from the remaining three (Figure 2C). These results also align with previous findings suggesting more consistent EV enrichment and sample preparation with the UF96 protocol. Differential protein analysis comparing UF96 and UC identified a substantial number of proteins with significantly different abundance, as represented in the report’s interactive volcano plot (Figure 3B) with accompanying tables. This graphic representation allows the user to explore the data and get immediate insights such as the observation that transmembrane proteins are present at higher levels using the UF96 approach. Proteins that were uniquely identified in either UF96 or UC are listed separately, in the “absent-from-DE” section.

**Figure 2.**
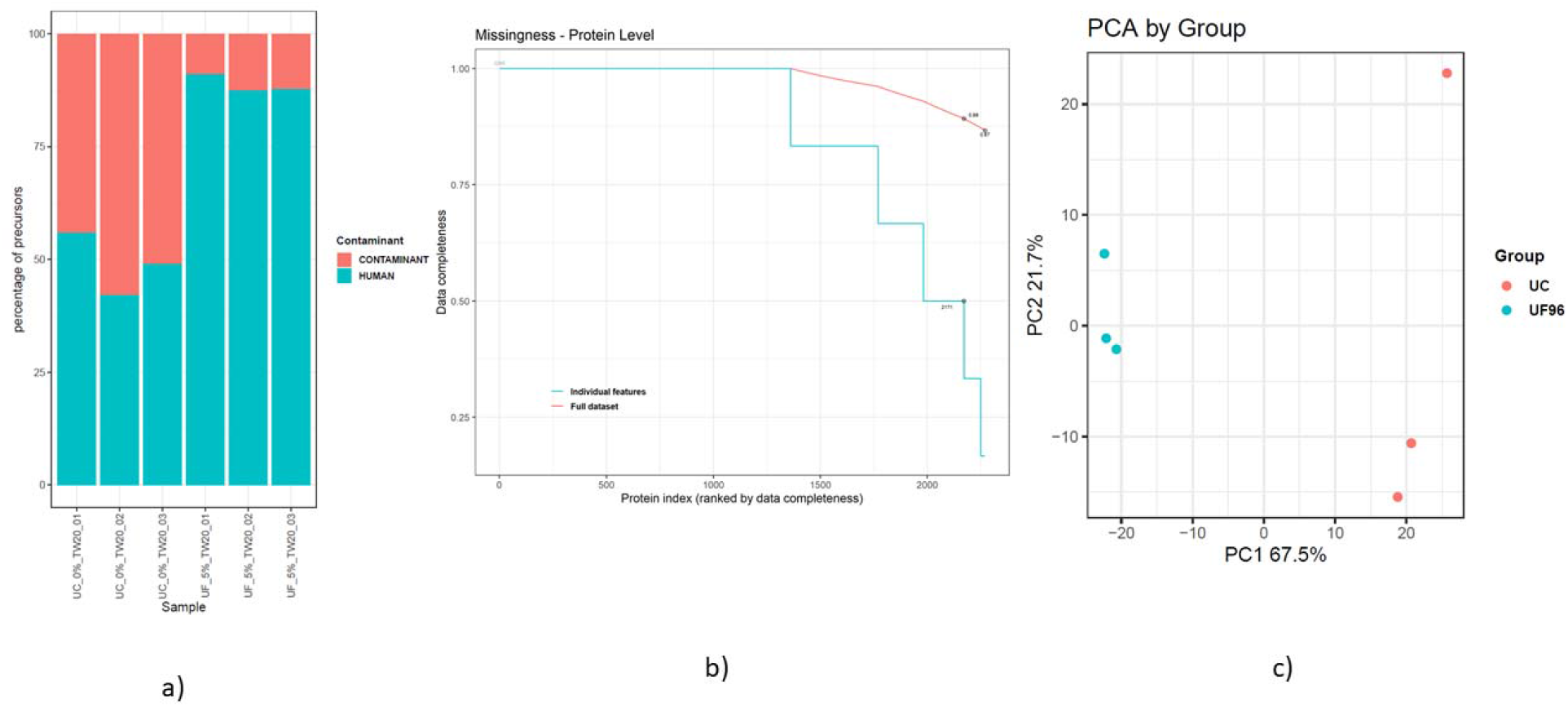
Quality control visualizations from the DiaReport EV module: (a) Proportion of identified precursors categorized by source (e.g., human vs. common contaminants). (b) Data completeness: proteins ranked by their data completeness across all samples. (c) Principal Component Analysis (PCA) of the experimental cohort; samples are colored by group (UF and UP98).

**Figure 3.**
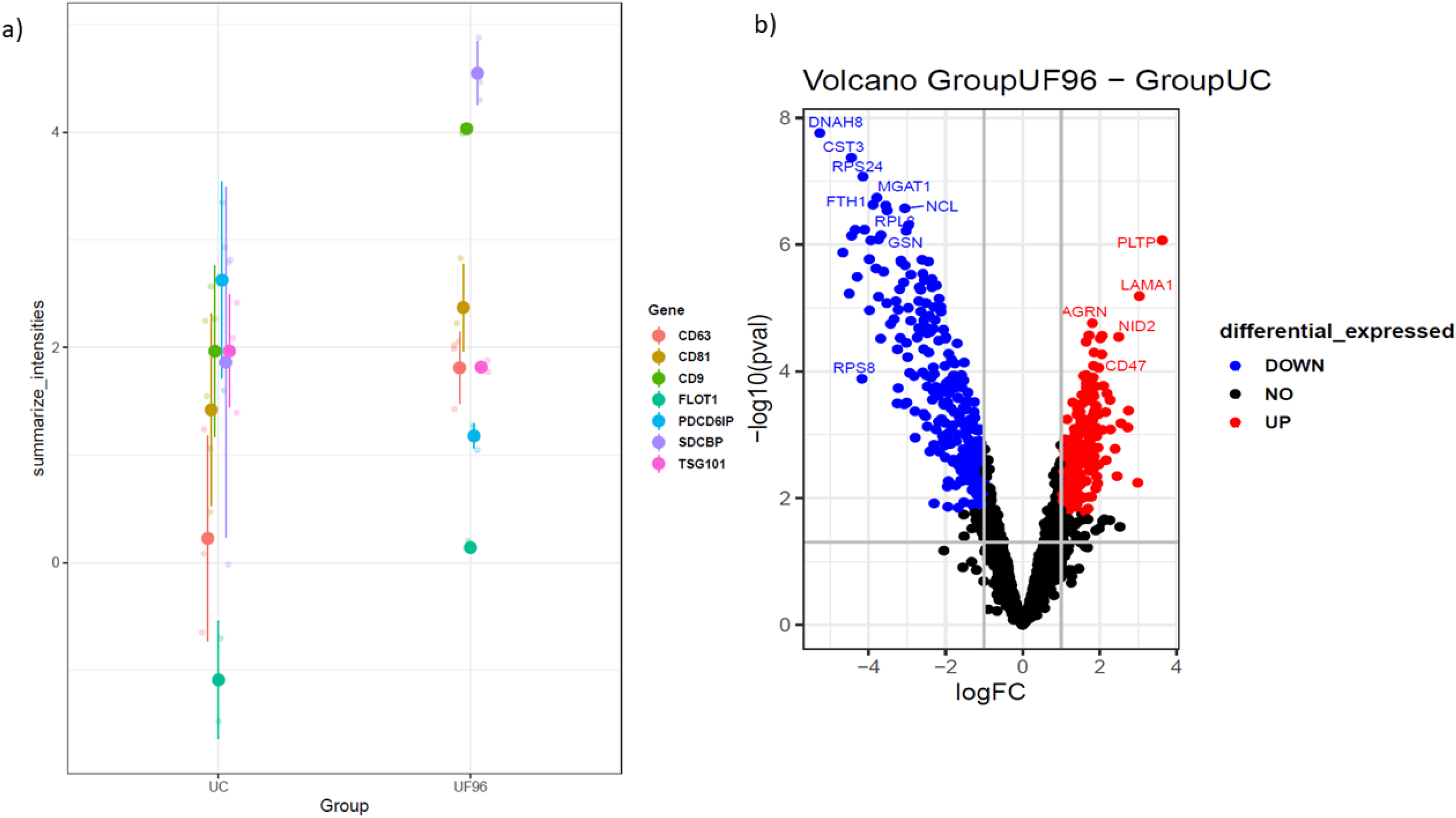
Visualizations from the DiaReport EV module: (a) Intensities of EV protein markers across the two experimental groups. Dark points represent the mean intensity for each marker, with error bars indicating the interval μ±σ. (b) Volcano plot of differentially expressed proteins, annotated by gene name (adjusted P-value <0.005 and ∣log2FC∣>1).

DiaReport provides the essential infrastructure to move from raw DIA-NN output to biological interpretation within a single, reproducible framework. For typical DIA clinical proteomics cohorts (50-200 samples), both the statistical analysis and report generation are complete within practical runtimes on a standard laptop (Supplementary Table S1). While the current implementation focuses on DIA-NN and MSqRob, the modular design facilitates future extensions to additional quantitative workflows and downstream analyses.

## Acknowledgements

We would like to thank the VIB Proteomics Core for validating and testing DiaReport, as well as Prof. Dr. Lieven Clement and the StatOmics group (UGent) for their technical support. An AI-based large language model was used for grammatical and linguistic refinement during the preparation of this manuscript

## Supplementary Information

**Supplementary Table S1. Indicative runtimes for DiaReport across different cohort sizes and storage configurations.**

The table reports wall-clock runtimes for differential expression analysis and HTML report generation using DiaReport on a standard laptop (32 GB RAM, Intel i7 processor). Measurements are shown for both precursor-and protein-level workflows, varying numbers of samples and contrasts, and execution from local storage or a network-mounted shared drive. “Analysis” corresponds to data processing and statistical modelling, while “Report” refers to Quarto-based HTML rendering. Total runtime is reported in seconds. Runtimes are intended as indicative values and may vary depending on hardware, storage performance, and dataset characteristics.

